# Longitudinal RNA Seq analyses reveal the prominent role of *Vagococcus* in broiler meat spoilage microbiome

**DOI:** 10.64898/2026.06.04.730080

**Authors:** Julia A. Manninen, Elio Nushi, Elina Jääskeläinen, Per Johansson, Johanna Björkroth

## Abstract

**Background:** Raw broiler meat products are highly perishable, and microbial activity is the main factor limiting their shelf lives. The spoilage microbiomes of broiler meat products have been studied mainly using traditional culturing methods and 16S rRNA gene amplicon sequencing, neither of which can show the activity of whole microbiome. Previous metatranscriptomic research of broiler spoilage has also remained limited to specific spoilage organisms rather than entire microbiomes.

**Results:** Our longitudinal study of broiler meat spoilage discovered the successions of active bacterial microbiomes and metabolic pathway activities of spoilers at 4°C and 6°C. Samples taken daily were subjected to metatranscriptomic analyses in combination with non-targeted metabolomic, traditional microbiology, and sensory analyses until advanced spoilage took place.

*Carnobacterium divergens*, *Carnobacterium maltaromaticum* and *Vagococcus proximus* were the most active species at both temperatures. Carnobacteria are known poultry spoilers whereas *Vagococcus proximus*, a species recently described, played an unexpected active role in the microbiome. It became dominant in samples stored at 6°C and its activity increased also at 4°C after the use-by date.

Central carbohydrate metabolism was the most common KEGG orthology pathway module of the microbiome at both temperatures. *C. divergens* and *C. maltaromaticum* showed stable metabolic profiles during the spoilage process, whereas *V. proximus* displayed a shift from high ATP synthesis activity to increased fatty acid and carbohydrate metabolism when spoilage advanced in samples stored at 4°C.

Non-targeted metabolomics showed similar metabolomic trends across both temperatures. At 6°C, time-dependent changes were generally more pronounced, and the spoilage markers tyramine and spermidine showed greater accumulation.

**Conclusions:** As expected, the rate of spoilage is higher at 6 than 4°C, however we did not anticipate a similar overall trajectory of the spoilage processes. Our results link *V. proximus* as a key active spoiler in broiler meat and demonstrate the efficacy of using RNA-seq together with metabolomics to decode the function of a meat spoilage microbiome. This demonstrates that spoilage microbiomes consist of active species we have been neglecting due to the technological limitations of the standard methods. Future studies targeting to the metabolism and detecting of *Vagococcus* are warranted.

## 1. Background

Broiler meat is a highly perishable food item that is rising in popularity around the world [1]. It is refrigerated and typically packaged under modified atmosphere (MA) to extend its shelf life and to control pathogenic bacteria. The MAs used may vary between countries in terms of oxygen content, but they typically contain carbon dioxide to suppress the growth of aerobic fast-growing spoilers [2–4]. Over the years, *Brochotrix, Carnobacterium,* and other psychrotrophic lactic acid bacteria (LAB)*, Photobacterium* and *Pseudomonas*, have been associated with the spoilage of MA-packaged broiler meat based either on traditional culturing or culture-independent analysis [5–10]. However, little is known about the interplay of these microbes within spoilage microbiomes during shelf life.

Spoilage microbiomes have mainly been assessed using 16S rRNA gene amplicon sequencing, which can confidently identify bacteria at the genus level. We chose to conduct a longitudinal metatranscriptomic study on MA packaged broiler meat and complemented it with metabolic, culture-based, and sensory analyses. Relatively few metatranscriptomic studies focusing on food spoilage microbiomes have been published. A longitudinal metatranscriptomic analysis of beef spoilage microbiome was performed by Hultman *et al.* [11] and Höll *et al.* [6] studied the metabolism of *Brochotrix thermosphacta* and *Carnobacterium divergens* in modified atmosphere packaged poultry using metatranscriptomics.

Compared to DNA-based methods, such as metagenomics, transcriptome analyses have the advantage of detecting living and active species and their functions. The aims of our research were to (I) investigate the succession of active bacterial species in broiler meat in the course of time, (II) identify the most prominent species at different timepoints, (III) observe pathway activities and study how metabolomic analyses are related to them, and (IV) identify changes in the pathway activity distribution in the course time while spoilage advanced. In addition, we studied how storage temperatures of 4°C and 6°C and lot-to-lot variation of meat was reflected in the spoilage microbiome development.

## 2. Materials and Methods

### 2.1 Broiler samples and longitudinal RNA sampling

Fresh skinless broiler leg meat cuts packaged in retail packages under modified atmosphere containing 30-40% CO_2_ and 70-60% N_2_ were obtained directly from a large-scale commercial producer. The packages originated from two separate production lots, one stored at 4°C and the other at 6°C throughout the shelf life and beyond until advanced spoilage took place.

From the packages stored at 4°C RNA was sampled every day starting from day 5 until day 14 after packaging. The final sampling point was 2 days after the use-by date. From the packages stored at 6°C, RNA was sampled daily from day 6 to day 10 after packaging, where day 10 was the use-by date. The shorter RNA collection window at 6°C was due to the faster spoilage rate. Three biological replicates with good RNA yield were sequenced for each sampling day.

### 2.2 Sensory evaluation

Immediately after RNA sampling, the sensory quality of meat was evaluated by 3-4 trained evaluators on a scale from 5 to 1 as described by Hultman et al. (2020). Sensory analysis was performed until the smell scores dropped to 1, meaning that the broiler meat was extensively spoiled. The quality attributes were considered to represent three stages based on the sensory scores and comments of the evaluators: acceptable product, early spoilage and late spoilage. Meat with a sensory score of 3 or above was classified as acceptable, between 3 and 1.75 as early spoiled and below 1.75 as late spoiled.

### 2.3 Microbiological analyses

Twenty grams of broiler meat was taken from each package and diluted 1:10 in peptone water (0.1 % peptone, 0.85 % NaCl) in a filter bag and homogenised using a Stomacher lab blender (Seward Ltd., Worthing, UK) at normal speed for 1 minute. A small aliquot of the liquid extracted from broiler was plated on four different media each day to estimate the bacterial concentration of different bacteria in the samples. The media used were Man-Rogosa-Sharpe (MRS) agar (Thermo Fischer), Plate count agar (PCA) (Thermo Fischer), Streptomycin thallous acetate actidione (STAA) agar (Thermo Fischer), and Violet red bile glucose (VRBG) agar (Thermo Fischer). Before counting the colonies, the VRBG plates were incubated at 37°C for 24 hours, the STAA plates at 25°C for 48 hours, the PCA plates at 30°C for 72 hours, and the MRS plates anaerobically at 25°C for 5 days.

### 2.4 RNA extraction and sequencing

A sample of 50g of broiler meat was mixed with 7.5ml of RNAlater (Thermo Fischer) and 2.5ml of peptone water, and homogenised using a Stomacher lab blender (Seward Ltd., Worthing, UK) at low speed for 30 s. The resulting mix was first centrifuged 3 min at 200 x *g* to remove the broiler tissue material. One millilitre of the resulting supernatant was pipetted and further centrifuged for 3 min at 13000 x *g* to pellet the bacterial cells. Then 300 µl of RNAlater was added to the pellet, and the resulting mix was stored at -70°C until RNA extraction.

RNA extraction was done using a Macherey-Nagel NucleoSpin RNA kit with a modified protocol. Special attention was paid to keeping the laboratory environment RNase-free, including wiping all surfaces and the centrifuge with RNase before starting. Cells stored in RNAlater were thawed and pelleted by centrifuging them for 10min at 16 000 x *g* at 4°C. RNAlater was decanted off and 700 µl of RA1 buffer was added on to the cell pellets. The buffer and cells were mixed by briefly vortexing before being transferred to Lysing Matrix E tubes that were kept on ice. The tubes were mixed in a bead beater (FastPrep) for 40 s at 5.5 m/s and before the tubes were again put on ice. After this, 500 µl of phenol-chloroform-isoamyl alcohol (25:24:1) was added to the tubes, which were vortexed for 30 s. The tubes were incubated at room temperature for 10 min and centrifuged for 10 min at 16 000 x *g* at 4°C. After centrifugation, 650 µl of the upper layer was pipetted into new 1.5ml tubes containing 650 µl chloroform. These new tubes were vortexed briefly and stored on ice for approximately 3 min. The tubes were centrifuged for 10 min at 16 000 x *g* at 4°C before 500 µl of the upper layer was pipetted into new 1.5ml tubes containing 5 µl betamercaptoethanol. Obtaining the lysate filtrate and the consequent steps were done according to the NucleoSpin RNA kit protocol, except the amount of 70% ethanol used was 480 µl.

Sequencing of the RNA samples was done by the Institute of Biotechnology of the University of Helsinki. The sequencing library was prepared with Illumina TruSeq and ribosomal depletion of the samples was done with chicken and bacteria riboPOOLS. Samples taken at 4°C were sequenced unpaired with Illumina NextSeq with 75bp sequencing length. Samples taken at 6°C were paired-end sequenced with Illumina NextSeq with 74bp forward reads and 88bp reverse reads.

### 2.5 Data processing pipeline

To generate pathway-level temporal activity and metatranscriptome profiles from raw RNA-seq data, a multi-stage processing pipeline composed of four hierarchical levels (Level 0–Level 4) was implemented. The pipeline follows the structure shown in Figure 1.

**Figure 1.**
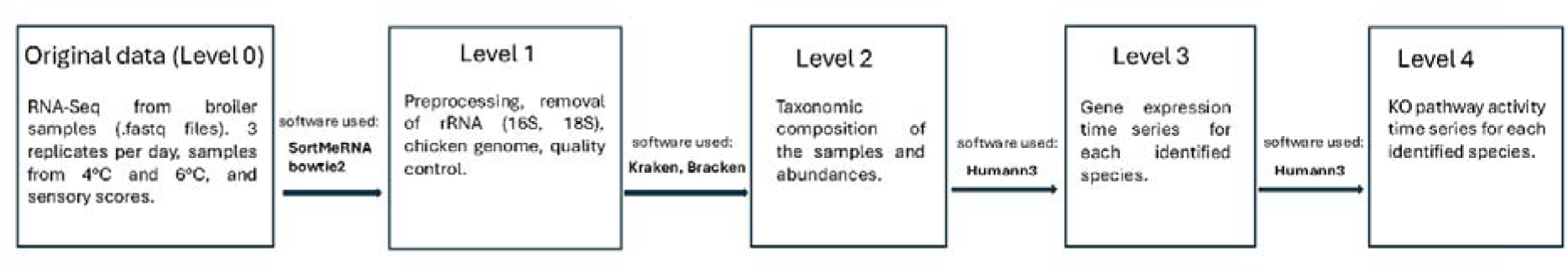
Metatranscriptomics pipeline used in this paper.

#### 2.5.1 Level 0 — Original Data (Raw RNA-seq Reads)

The starting point of the analysis consists of raw RNA-seq reads in the form of fastq files obtained from the sequencing of the broiler RNA samples collected during the experiments. For each day, three biological replicates were sequenced. These raw reads represent the complete microbial and host transcriptional output prior to any processing.

#### 2.5.2 Level 1 – Preprocessing of the data

Raw RNA sequences were first processed with SortMeRNA [12] to remove any remaining 16S and 18S rRNA from the reads. After this, the data was processed for species abundance analysis using a protocol originally published by Lu *et al.* [13]. Removing the host RNA was done by aligning the sequences to *Gallus gallus* (chicken) with Bowtie2 v. 2.5.3 [14]. NCBI accession GCA_016699485.1 (BioProject PRJNA660757) was indexed with bowtie2-build and used as reference index.

#### 2.5.3 Level 2 — Taxonomic Composition (Kraken2 and Bracken)

Kraken2 v. 2.1.5 [15] was used to estimate the abundances of bacteria and other micro-organisms in the RNA samples. A database for Kraken2 was built by downloading the bacteria, fungi and viral databases of Kraken2 with kraken2-build. The database was complemented with some lacking spoilers i.e. *Dellaglioa algida* (GCF_030127605), *Dellaglioa carnosa* (GCF_026948345), *Pseudolactococcus carnosus* (GCF_006770265), *Photobacterium carnosum* (GCF_021236965), *Vagococcus fessus* (GCF_003987565), and *Vagococcus proximus* (GCF_029143285). Their fasta sequences were downloaded from the NCBI database FTP and added in with the other databases before the final Kraken2 database was built.

The results from Kraken2 were used to run Bracken [16] to refine the bacterial species abundances, as Bracken corrects biases in k-mer based assignments. The k-mer database for Bracken was built specifically for the read length, which was 75bp for the 4°C samples and 74bp for 6°C. This step yielded the list of microbial species detected in each sample, and the species-level abundance profiles across all time points. These taxonomic profiles provided the basis for species-resolved functional analyses. Results from this step were also visualised with Krona [17] to provide an overview of the succession of active species in the spoilage microbiome.

#### 2.5.4 Level 3 — Gene Expression Time Series (HUMAnN)

In the third level, functional gene families were reconstructed for each identified species using HUMAnN [18]. HUMAnN maps reads to species-specific pangenomes, producing a gene-family expression profile for each species across time. The output of this step was gene expression time series per species, enabling downstream modelling of temporal gene-level activity. Conventionally HUMAnN uses MetaPhlAn [19] for performing taxonomic profiling; in our setup, however, we skipped this step and instead used the taxonomic profiling obtained from running Bracken as a best practice suggested by Pereira-Marques *et al.* [20].

#### 2.5.5 Level 4 — Pathway Activity Time Series (HUMAnN)

At the final processing level, HUMAnN converted gene-family abundances into KEGG Orthology (KO) pathway activity profiles. For each species, gene families were mapped to KO identifiers and aggregated into pathway modules. This produced a species-resolved KO pathway activity time series, representing the temporal trajectory of key metabolic functions in the community.

### 2.6 Non-targeted metabolomics

#### 2.6.1 Sample preparation

Meat samples were stored at −80°C until analysis. Prior to extraction, frozen samples were thawed at 8°C for two hours and then approx. 100 mg was weighed into homogeniser tubes. For metabolite extraction, cold methanol (80% v/v) was added in a ratio of 400 µl per 100 mg of sample. The samples were homogenised using metal beads at 6 m/s for 30 s. The samples were then kept on ice for 15 min, vortexed for 10 s at room temperature, and centrifuged at 17 000 x *g* at 4 °C for 10 min. After centrifugation, the samples were transferred to 96-well filter plates. Filtering was performed by centrifuging the plate at 700 x *g* at 4 °C for 5 min. The quality control (QC) sample was prepared by collecting 50 µl from each sample and pooling the aliquots into four wells. A blank sample and a sample containing only 80% methanol were prepared.

#### 2.6.2 LC–MS analysis

The samples were analysed by liquid chromatography–mass spectrometry using a 1290 Infinity II UHPLC system (Agilent Technologies, Santa Clara, USA) coupled to a high-resolution QTOF mass spectrometer (Agilent 6546 with Jet Stream ion source, Agilent Technologies). All sample handling, preparation, and LC–MS analyses were performed at Afekta Oy (Kuopio, Finland). The analytical method has been described in more detail by Hanhineva *et al.* [21] and Klåvus *et al.* [22]. In brief, a Zorbax Eclipse XDB-C18 column (2.1 × 100 mm, 1.8 µm; Agilent Technologies) was used for the reversed-phase (RP) separation and an Acquity UPLC BEH amide column (Waters) for the hydrophilic interaction liquid chromatography (HILIC) separation. After each chromatographic run, the ionisation was carried out using jet stream electrospray ionisation (ESI) in the positive and negative mode, yielding four data files per sample. The collision energies for the MS/MS analysis were selected as 10, 20, and 40 V, for compatibility with spectral databases.

#### 2.6.3 Data analysis

Peak detection and alignment were performed in MS-DIAL ver. 4.90 [23]. For the peak collection, m/z values between 50 and 1500 and all retention times were considered. The amplitude of minimum peak height was set at 3000. The peaks were detected using the linear weighted moving average algorithm. For the alignment of the peaks across samples, the retention time tolerance was 0.05 min and the m/z tolerance was 0.015 Da. The solvent background was removed using solvent blank samples under the condition that, to be kept for further data analysis, the maximum signal abundance across the samples had to be at least five times that of the average in the solvent blank samples.

After the peak picking, a total of 42,138 molecular features were included in the data preprocessing and clean-up step. Low-quality features were flagged and discarded from statistical analyses. Molecular features were only kept if they met all the following quality metrics: a low number of missing values (present in more than 70% of the QC samples, present in at least 50% of samples in at least one study group), a relative standard deviation (RSD) below 20%, a D-ratio below 10%. In addition, if either the RSD or D-ratio was above the threshold, the features were kept if their classic RSD and basic D-ratio were all below 10%. The signals were normalised for signal drift and batch effect. Missing values were imputed using Random Forest imputation for high-quality features and using simple imputation with a value of 0 for low-quality features.

#### 2.6.4 Compound identification

The chromatographic and mass spectrometric characteristics (retention time, exact mass, and MS/MS spectra) of the selected molecular features were compared with entries in an in-house standard library and publicly available databases, such as METLIN and Human Metabolome Database (HMDB), as well as with published literature. The annotation of each metabolite and the level of identification was given based on the recommendations published by the Chemical Analysis Working Group (CAWG) Metabolomics Standards Initiative (MSI) [24].

## 3. Results

### 3.1 Sensory scores and microbiological results

According to the evaluators, smell deteriorated faster than appearance at both 4°C and 6°C. The average smell scores at 4°C and 6°C fell to below 3 on days 11 and 8, respectively. Appearance degraded more slowly, with average appearance scores falling to below 3 on day 13 and 9, respectively (Supplementary Figure 1).

The final concentration of bacteria on MRS and PCA was similar at both 4°C and 6°C, reaching over 10^8^ CFU/g by the end of the experiments. At 4°C, STAA reached a bacterial concentration of between 10^5^ and 10^6^ CFU/g and VRBG 10^5^ CFU/g. At 6°C, the VRBG count was higher, reaching a concentration of between 10^6^ and 10^7^ CFU/g, whereas STAA was lower at around 10^4^ CFU/g (Supplementary Figure 2). The bacterial communities were in the stationary phase for both early and late spoilage at both 4°C and 6°C, whereas the sensory scores fell faster. At 6°C, the broiler samples were already in late spoilage by the use-by date, whereas at 4°C, the samples were in early spoilage at the time of the use-by date (Supplementary Figure 3).

### 3.2 Experiment at 4°C

#### 3.2.1 Species succession

At 4°C a clear succession of active species in the spoilage microbiome was detected. In the beginning of the experiment, a wide variety of bacteria along with some fungal species were active. LAB comprised a relatively small percentage (about 22%) of them, whereas *Propionibacterium* and gram-negative genera, such as *Rahnella*, *Pseudomonas,* and *Yersinia,* accounted for a large share. Many of the microbes active in the first samples were not associated with early or late spoilage microbiomes, showing how the initial contaminating community was marginalised during the succession (Figure 2).

**Figure 2.**
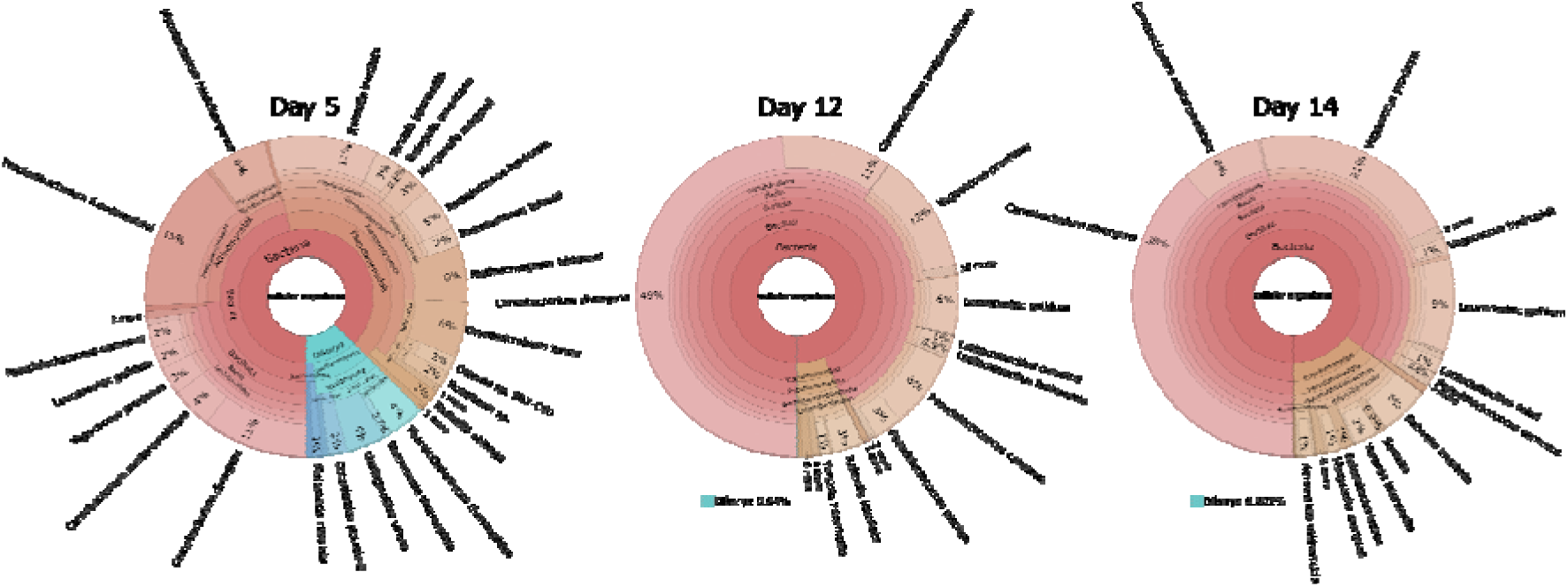
Succession of the active spoilage microbiome at 4°C visualised with Krona [17].

The share of gram-negative species in the active microbiome decreased as the broiler became closer to its use-by date. In the course of time, LAB became increasingly dominant in the active microbiome. By the use-by date, they comprised over 90% of the active microbiome. The most prominent species in the active microbiome were *Carnobacterium divergens, Carnobacterium maltaromaticum*, *Vagococcus proximus,* and *Leuconostoc gelidum* (Figure 2)*. C. divergens* especially had the highest read count of all species in the samples (Figure 3). Towards the end of the shelf-life, carnobacteria accounted for over half of the RNA reads in the samples. After passing the use-by date, the major changes related to the active microbiome were the increasing share of vagococci (Figures 2 and 3) and the slight increase in the abundance of gram-negative species in the microbiome. Reads aligning to fungal species also fell quickly after day 5, with their proportion of the microbiome falling to below 1% by day 8 (Figure 2).

**Figure 3.**
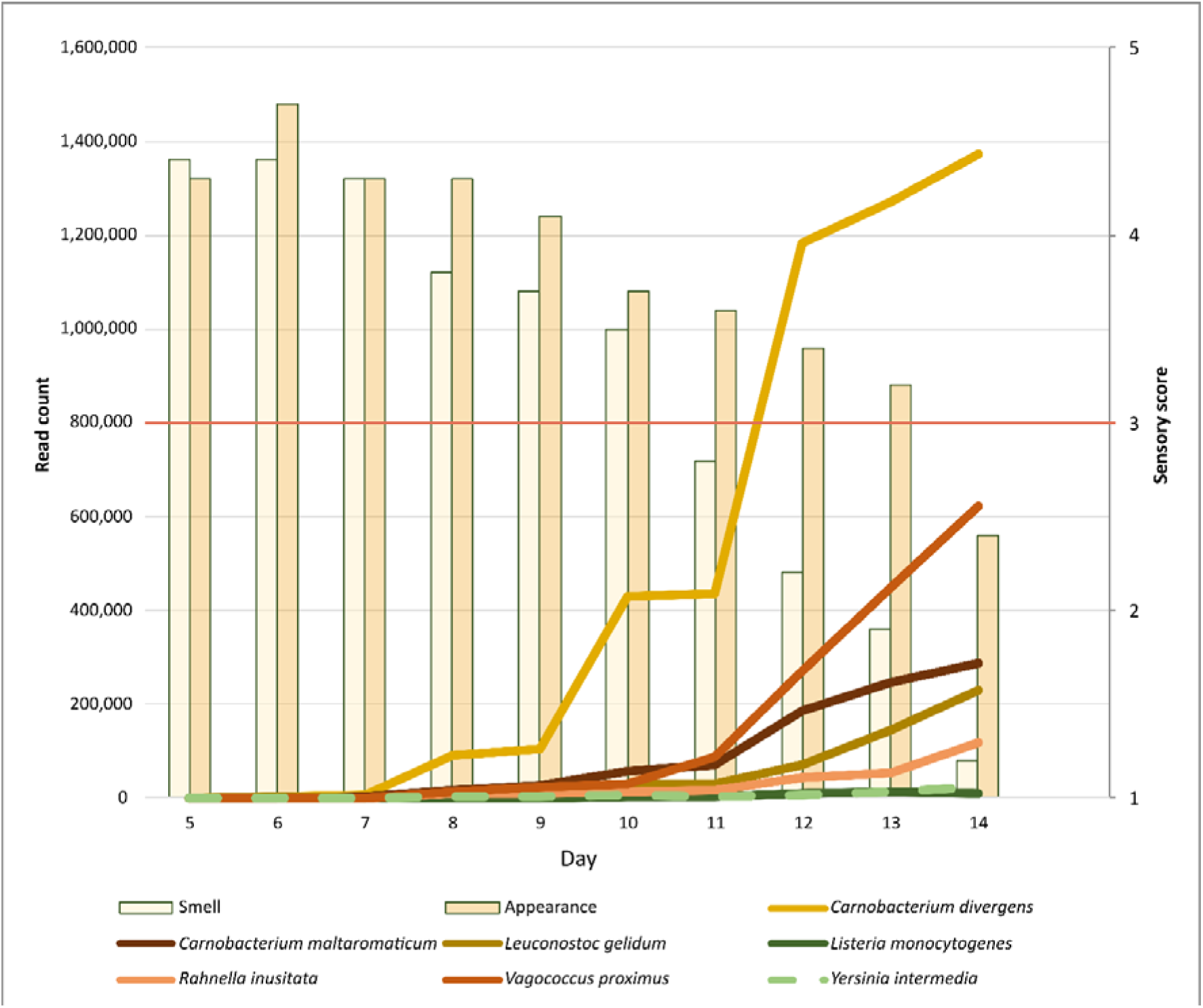
Average read counts of few selected bacteria in the active spoilage microbiome at 4°C.

#### 3.2.2 Pathway activity

The most active KEGG module in the spoilage microbiome was central carbohydrate metabolism, encompassing metabolic pathways such as glycolysis, citrate cycle and pentose phosphate pathway. ATP synthesis included pathways for many enzymes related to energy metabolism. Carbon fixation mainly consisted of pathways related to metabolic cycles, such as the Calvin cycle, Arnon-Buchanan cycle, as well as the production of acetate. Other carbohydrate metabolism encompassed pathways such as nucleotide sugar biosynthesis and ascorbate degradation.

*C. maltaromaticum* and *C. divergens* had a similar distribution of KEGG modules across the different phases of spoilage (Figure 4). Their pathway distributions mirrored the distribution of the whole spoilage microbiome, as they made up most of the active spoilage microbiome (Figure 2).

**Figure 4.**
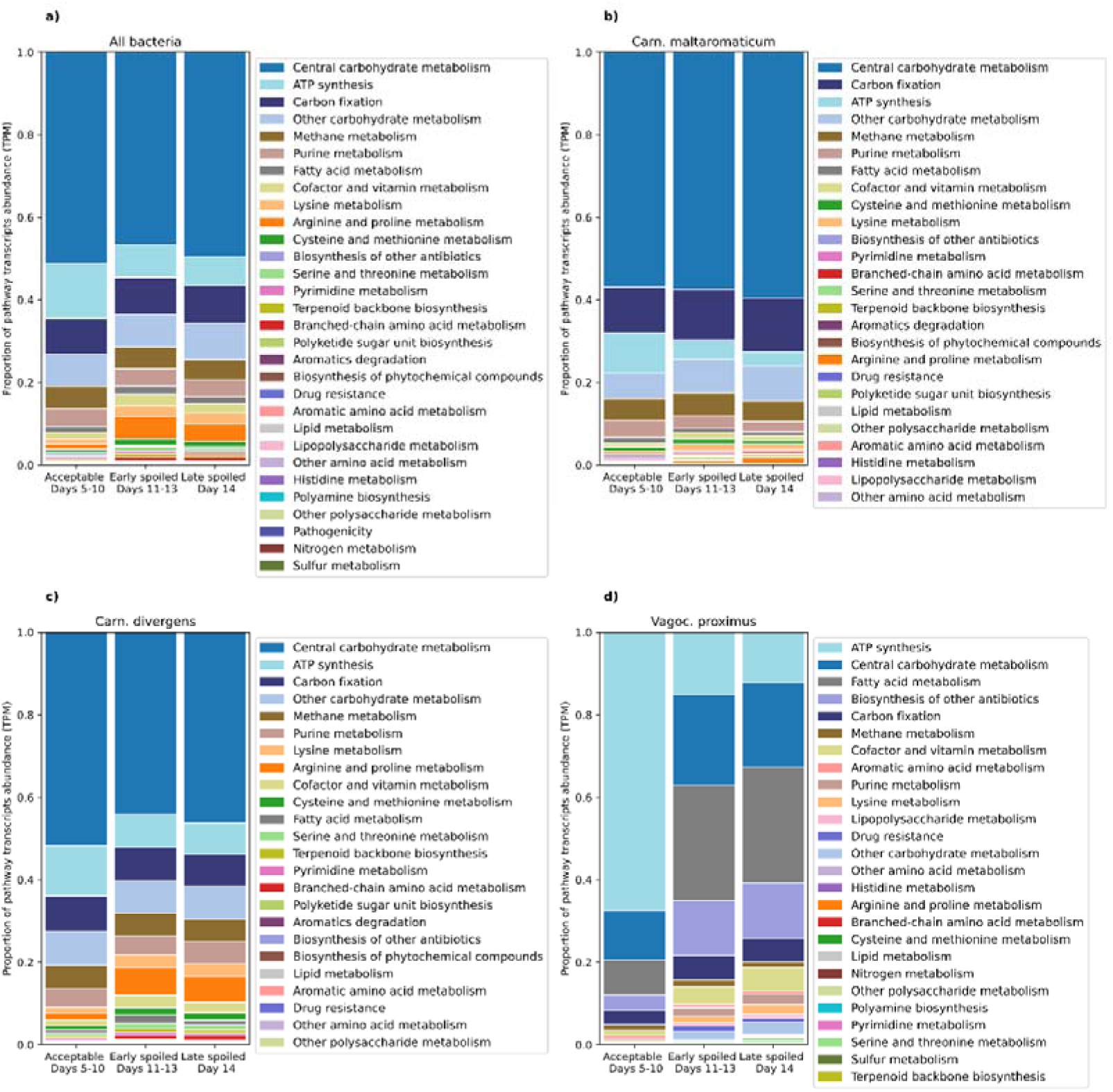
Distribution of metabolic pathway activities of the whole microbiome, *C. maltaromaticum, C. divergens,* and *V. proximus* during the spoilage at 4°C.

*V. proximus* had a different distribution of pathway activity from the other dominant species in spoilage microbiome. It had highly active ATP synthesis in the acceptable product phase and a noticeable change in the activity distribution across the different spoilage phases. ATP synthesis fell after the acceptable product phase and carbohydrate and fatty acid metabolisms rose as spoilage advanced (Figure 4). The metabolic pathway module classified under antibiotic production was annotated as aurachin A biosynthesis by KEGG. The fatty acid metabolism category consisted mainly of meromycolic acid biosynthesis, and fatty acid biosynthesis initiation and elongation.

### 3.3 Experiment at 6°C

#### 3.3.1 Species succession

At 6°C, the succession of active species was, as expected, faster than at 4°C. By the time the first RNA sample was obtained on day 6, the succession had advanced well, and the microbiome resembled the 4-degree microbiome near the use-by date. At 6°C LAB were clearly the dominant order of bacteria, with gram-negative bacteria making up only a small portion of the active microbiome. *V. proximus* was the most dominating species in the active spoilage microbiome, with *C. divergens* and *C. maltaromaticum* also showing noticeable shares (Figure 5 and Supplementary Figure 4).

**Figure 5.**
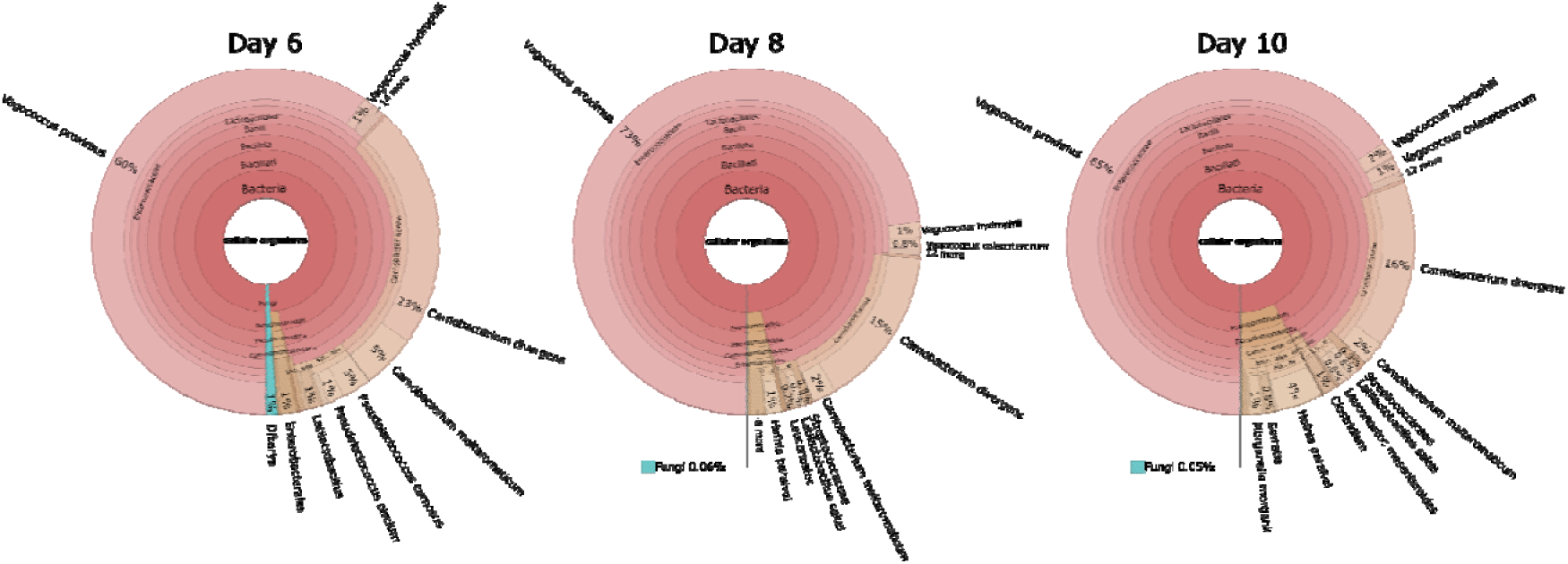
Succession of the active spoilage microbiome at 6°C with Krona [17].

#### 3.3.2 Pathway activity distribution

Pathway activity at 6°C was dominated by different modules of carbohydrate and carbon metabolism. As with the experiment at 4°C, *V. proximus* had a different pathway activity distribution from the other dominant bacterial species with highly active fatty acid metabolism, biosynthesis of antibiotics, and ATP synthesis. *C. maltaromaticum* was far less abundant at 6°C than at 4°C, only crossing the 5% abundancy threshold during the acceptable product phase (Figure 6).

**Figure 6.**
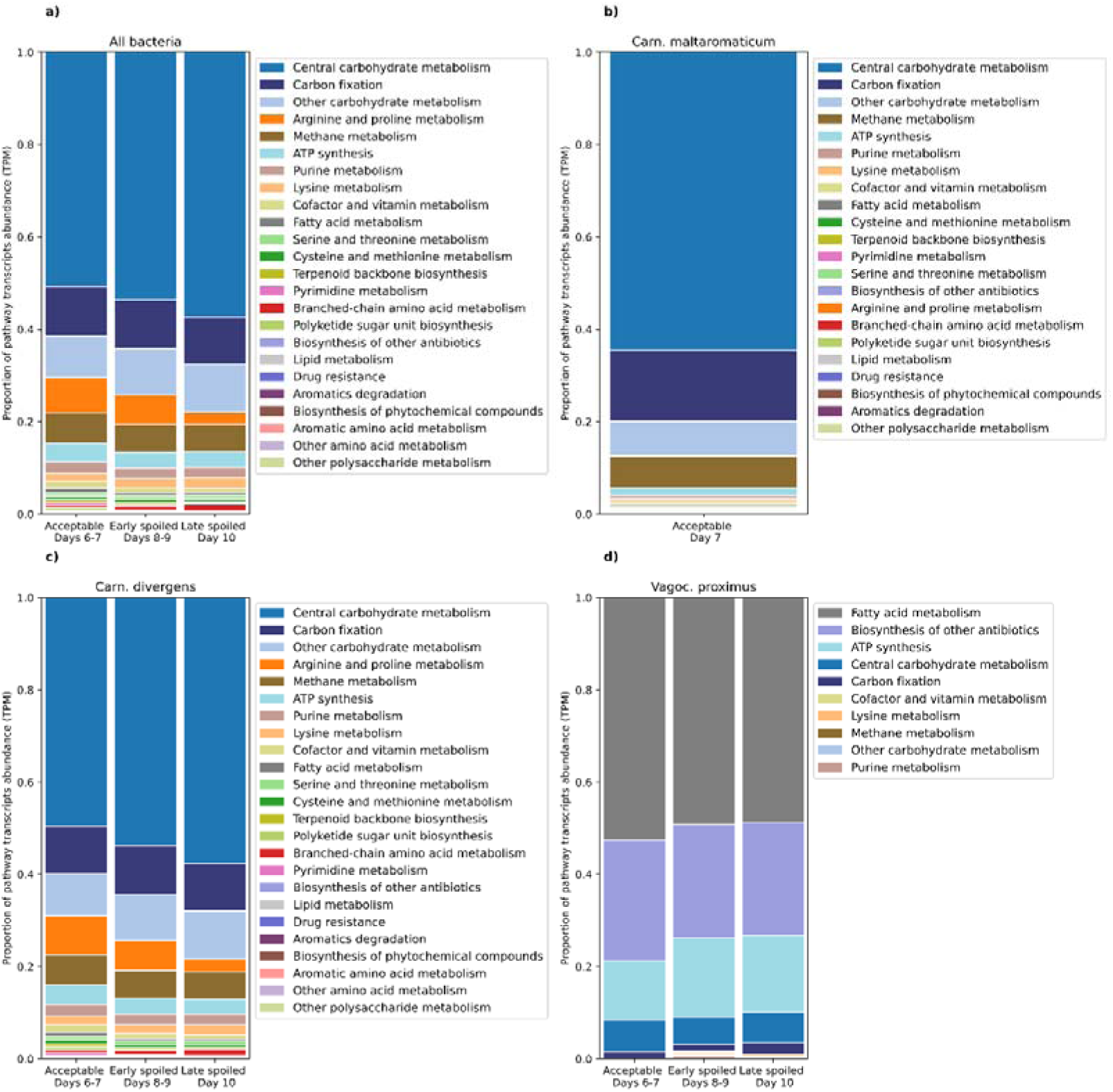
Distribution of metabolic pathway activities of the whole microbiome, *C. maltaromaticum, C. divergens,* and *V. proximus* during the spoilage at 6°C.

### 3.4 Metabolomic profiling

The non-targeted metabolomic analysis of broiler samples using RP chromatography and HILIC in positive and negative ionisation modes yielded 31,712 molecular features after peak picking, drift correction, and removal of low-quality signals. At the molecular feature level, more than half of the features were associated (Spearman correlation, q < 0.05) with the sensory evaluation score (18,438 features out of 31,712) and the bacterial growth stage (15,616 features) (Figure 7). Most of these features (14,653) are common between both variables. A total of 10,930 features were associated with time, whereas only 29 features were associated with temperature. When both time and temperature were included in the statistical model as an interaction, there were 433 significantly associated features.

**Figure 7.**
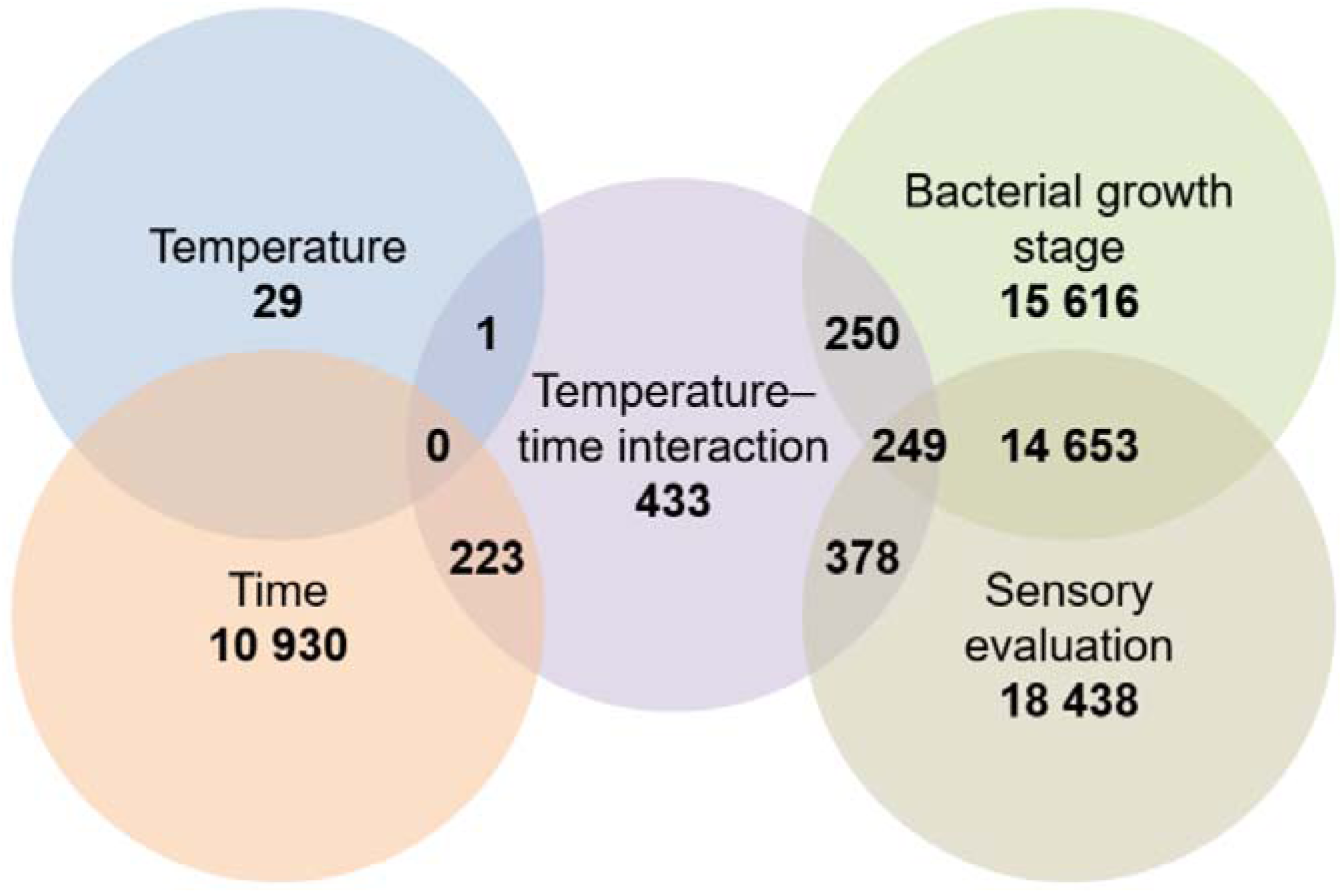
Euler diagram showing the number of good-quality molecular features (out of a total of 31,712) that have a q-value < 0.05 in the different statistical comparisons.

Among the annotated metabolites (MSI level 2–3), several compounds displayed clear time-dependent trends (Table 1). Spermidine increased over time at 6°C, while remaining relatively stable at 4°C. Tyramine showed pronounced accumulation, particularly at 6°C, consistent with microbial decarboxylation of amino acids. Proline-Glycine-Proline (Pro-Gly-Pro) levels increased over time at both storage temperatures, with slightly higher levels observed at 6°C.

**Table 1.** Longitudinal metabolite profiles of broiler meat stored at 4°C and 6°C. Mean peak areas (×1,000) of three replicate samples per time point for selected metabolites. Peak areas correspond to the relative abundance of each compound measured by non-targeted metabolomics using RP and HILIC chromatography.

| Compounds | 4°C |  |  |  |  |  |  | 6°C |  |  |  |
| --- | --- | --- | --- | --- | --- | --- | --- | --- | --- | --- | --- |
|  | Day<br>1 | 5 | 8 | 9 | 11 | 12 | 13 | 1 | 5 | 8 | 10 |
| **Nucleotide degradation products** |  |  |  |  |  |  |  |  |  |  |  |
| IMP | 6719,4 | 4649,8 | 3474,2 | 2943,1 | 2074,4 | 2281,1 | 2337,4 | 4949,6 | 3012 | 1359 | 243,9 |
| Inosine | 19,8 | 24,7 | 23,8 | 27,2 | 24,3 | 24,8 | 23,2 | 22,6 | 26,6 | 16,9 | 3,5 |
| Hypoxanthine | 6,3 | 10,2 | 13,7 | 13,4 | 15 | 16,7 | 18,7 | 9,5 | 14,3 | 23,2 | 27,5 |
| AMP | 4,1 | 28,4 | 44,5 | 64,2 | 53,6 | 78,1 | 83,8 | 5,8 | 47,4 | 111,5 | 84,7 |
| Xanthine | 448,7 | 680,7 | 562,7 | 507,8 | 810,2 | 845,6 | 695,1 | 554,8 | 594,6 | 773,2 | 896,5 |
| Uric acid | 6,2 | 13 | 17,8 | 16,7 | 19,7 | 21,3 | 14,4 | 8,8 | 14,6 | 29,8 | 35,2 |
| **Biogenic amines** |  |  |  |  |  |  |  |  |  |  |  |
| Tyramine | 19,7 | 364,8 | 2996,8 | 4619 | 6010,7 | 6305 | 7675,3 | 14,5 | 1622,4 | 9116,5 | 9612,1 |
| Putrescine | 1178,7 | 1130,2 | 958,7 | 1142,7 | 1282,1 | 2094,1 | 2601,2 | 1268,7 | 1379,4 | 3339,5 | 8465,8 |
| Histamine | 597 | 514 | 709,8 | 535,7 | 659,1 | 766,8 | 1312,7 | 480,9 | 423,9 | 1873,1 | 6588,1 |
| Tryptamine | 0,9 | 3,7 | 14,7 | 17,3 | 260,9 | 195,9 | 1022 | 0,8 | 1,8 | 608,6 | 698,6 |
| Spermidine | 501,3 | 547,3 | 514,4 | 511,7 | 532,8 | 578,3 | 561,9 | 492,2 | 587,1 | 787,2 | 895,8 |
| **Free amino acids and related compounds** |  |  |  |  |  |  |  |  |  |  |  |
| Proline | 0,2 | 0,2 | 0,3 | 0,3 | 0,5 | 0,9 | 1,2 | 0,2 | 0,2 | 1,5 | 1,9 |
| Arginine | 12,4 | 15,1 | 17,1 | 16,7 | 16,2 | 15,9 | 14,3 | 13,5 | 16,2 | 11 | 2,9 |
| Citrulline | 1,5 | 1,8 | 2,2 | 2,1 | 3,2 | 4,1 | 4,8 | 1,8 | 2,1 | 4,8 | 3,8 |
| Ornithine | 0,7 | 0,9 | 1,2 | 0,9 | 2,1 | 4 | 5,5 | 0,6 | 0,9 | 7,5 | 9,4 |
| Lysine | 7235,8 | 10394 | 11688,5 | 11646,6 | 11421,6 | 12619,7 | 12635,4 | 8975,4 | 11838,6 | 12746,3 | 12735,4 |
| Tyrosine | 14653,1 | 23539,4 | 21545,4 | 17984,8 | 12882,5 | 15247,9 | 11503,6 | 20729,9 | 26089,6 | 4884,2 | 2545 |
| Phenylalanine | 349 | 333,4 | 271,3 | 294,9 | 289,6 | 307,5 | 310,1 | 360,2 | 333,6 | 328 | 313,1 |
| Tryptophan | 581,4 | 1255,9 | 2252,5 | 2223,5 | 2109,6 | 2907,9 | 3033 | 947 | 2052,9 | 3191,2 | 3339,6 |
| Asparagine | 3,1 | 4,5 | 5,1 | 5,3 | 5,2 | 6 | 6,1 | 4,4 | 5,8 | 6,5 | 6,1 |
| Histidine | 368,6 | 395,7 | 415,3 | 401,7 | 383,2 | 388 | 381,5 | 375,6 | 400,7 | 366,9 | 360 |
| **Peptides** |  |  |  |  |  |  |  |  |  |  |  |
| Pro-Gly-Pro | 46,5 | 213,3 | 332,6 | 353,9 | 402,1 | 492,9 | 532,6 | 93,8 | 359 | 634,9 | 725 |
| **Carbohydrate-related metabolites** |  |  |  |  |  |  |  |  |  |  |  |
| Glucose | 210,5 | 261,5 | 232,7 | 260,3 | 227,2 | 256,6 | 247,1 | 174 | 175,6 | 170,9 | 176,2 |
| Glucose-6-phosphate | 85,2 | 185 | 182,7 | 215,4 | 259 | 204,3 | 184 | 104,2 | 236,2 | 177,7 | 17,5 |
| Pyruvate | 34,7 | 52,8 | 37,9 | 39,4 | 23,5 | 24,6 | 18,9 | 36 | 50,1 | 7,3 | 5,3 |
| Ribose-5-phosphate | 1122,1 | 860,5 | 571,1 | 596 | 475,9 | 433,4 | 359,6 | 1103,4 | 715,9 | 269 | 75,9 |

| Redox-, energy- and cofactor-related metabolites |  |  |  |  |  |  |  |  |  |  |  |
| --- | --- | --- | --- | --- | --- | --- | --- | --- | --- | --- | --- |
| NADP+ | 7,8 | 12,4 | 10,8 | 7,4 | 4,4 | 3,8 | 2 | 11,8 | 7,5 | 1,3 | 0,9 |
| NADPH | 2,1 | 21,5 | 22,6 | 25,2 | 17,1 | 18,6 | 16,8 | 14,5 | 18,1 | 8,1 | 4,3 |
| Glutathione | 3851,1 | 2954,8 | 2222,1 | 1875,3 | 1909,1 | 1652 | 1760 | 3477,2 | 2205 | 1144,3 | 743,1 |
| Riboflavin | 47,2 | 101,1 | 46,5 | 76,4 | 81 | 95,3 | 98,4 | 57,4 | 51,5 | 94,3 | 164 |
| Flavin mononucleotide | 56,5 | 71 | 71,4 | 72,4 | 95,6 | 97,6 | 84,9 | 66,7 | 82,3 | 97,8 | 118,4 |
| Flavin adenine dinucleotide | 8,9 | 13,2 | 6,1 | 6,5 | 5,4 | 8 | 8 | 6,4 | 5,3 | 6,7 | 10,5 |
| Thiamine | 5577,3 | 8473,5 | 7677,4 | 8265,3 | 7919,4 | 9183,7 | 8567,6 | 7491,1 | 8449,4 | 9173,6 | 8999,3 |

Nucleotide degradation was reflected by a progressive decrease in inosine monophosphate (IMP) at both temperatures. Inosine remained relatively stable at 4°C but decreased at 6°C, whereas hypoxanthine increased over time under both storage conditions.

Most free amino acids showed moderate time-dependent increases over the storage period. The dynamics of certain amino acids, such as tyrosine, may reflect both accumulation and partial conversion into biogenic amines (e.g., tyramine).

At 4°C, changes in amino acids and nucleotide-related compounds were gradual: IMP decreased steadily, inosine levels remained fairly constant, and hypoxanthine increased slightly. At 6°C, time-dependent changes were generally more pronounced, with accelerated nucleotide degradation, declining inosine, increased hypoxanthine, and greater accumulation of biogenic amines.

Comparing the two temperatures, the primary difference lay in the rate of metabolic changes rather than in the emergence of distinct metabolic patterns. Overall trends were qualitatively similar, but the higher storage temperature accelerated nucleotide degradation, biogenic amine formation, and moderately amplified the accumulation of certain amino acids and peptide fragments.

## 4. Discussion

Despite coming from two different production lots and being stored at two different temperatures, the spoilage process in our experiments was remarkably similar. The three most dominant species were the same, just occurring in different proportions. Spoilage process at both temperatures also showed similar metabolic trends. While the changes occurred faster at 6°C, they followed the same general trajectory as at 4°C that was consistent with the succession of LAB and the shifts in pathway activity observed in the transcriptomic data. Pathway activity distribution of the whole spoilage microbiome was similar at both temperatures. This is likely due to the strong selective pressure MA packaging and particularly CO_2_, exerts on the spoilage microbiome of meat [25, 26]. Säde *et al.* [27] also showed that despite the initial differences in early bacterial communities of MA packaged meat, the communities rapidly became more similar due to succession when spoilage advanced.

The main difference between the two temperatures lay in the rate of spoilage. At 4°C, the broiler meat was in the early spoilage phase at its use-by date, whereas at 6°C it had already reached late spoilage by the time of the use-by date. For consumers, this means a substantial difference in quality, as broiler meat products in the late spoilage phase are very unpleasant due to strong odours and unappealing appearance. The metabolomic patterns observed, such as increased polyamines and biogenic amines, reflect these accelerated microbial processes. The higher temperature accelerated and intensified the accumulation of biogenic amines, which are known to cause unpleasant smells and to degrade the quality of meat products. The rising levels of spermidine in the 6°C samples likely contribute to the rapid deterioration of smell scores compared to the samples stored at 4°C, which had a stable and lower spermidine amount. In high concentrations biogenic amines can also be toxic [28], making the storage temperature a matter of food safety also from the metabolic point of view.

Spermidine is typically produced from putrescine but can also be produced from agmatine [29]. However, spermidine production is mainly associated with gram negative bacteria and streptococci and has not been detected in either vagococci or carnobacteria, and thus we cannot associate it with the active microbiome without further studies. Tyramine, another biogenic amine, can be produced by carnobacteria [30, 31]. *C. divergens* was predicted to be able to produce the biogenic amines cadaverine, tyramine and GABA *in situ* in broiler meat by Höll *et al.* [6]. The type strain of *V. proximus* does have a tyrosine decarboxylase gene (Uniprot accession A0ABT5X1I6), although there has not been any *in vitro* evidence that it is able to produce tyramine.

The two-degree difference in temperature affected the dominant species of the spoilage microbiome. *V. proximus*, a prominent bacterium at 4°C became strongly dominant at 6°C whereas *C. divergens* was less abundant (Supplementary Figure 5). This indicates either a better fitness of *V. proximus* at 6°C or lower fitness of *C. divergens*.

The pathway activity distributions indicate that *V. proximus* behaved differently than the other dominant spoilers during the spoilage process. At 6°C, *V. proximus* did not practically exhibit any central carbohydrate metabolism, whereas fatty acid metabolism, which mainly consists of biosynthesis of different lipids, was pronounced. At 4°C, *V. proximus* started with a high ATP synthesis activity, but then the pathway activity also begun to shift towards fatty acid metabolism as the spoilage advanced. This change in metabolic activity coupled with the growth of the microbial community suggests that *V. proximus* stopped growing by the end of the acceptable product phase and started to synthesise fatty acids or lipids. The changing metabolism at 4°C makes *V. proximus* a more likely culprit for the decreasing sensory values in those samples, as from the three dominant species in the samples it is the only one that has a significant change in its pathway activities between acceptable and spoiled products. *C. divergens* and *maltaromaticum* had stable pathway activities during the spoilage, with only minor changes in pathway activities.

Until now, little has been known about the role of vagococci in the spoilage of meat products. Lauritsen *et al.* [8] detected vagococci in whole broiler meat packaged under MA of 80% O_2_/20% CO_2_ and reported that they constituted a dominating part of the later shelf-life microbiota. *V. proximus* was described only recently in 2023 [32] from MA (30% CO_2_/70% N_2_) packaged broiler meat, but its role in spoilage remained unverified. A major obstacle for detecting vagococci from meat samples has been the lack of selective media, which could partly explain why it has not been widely recognised as a spoiler until now. Further studies should be done on the spoilage potential of vagococci to better understand their metabolism and role in meat spoilage.

Studies from other countries often detect high levels of a well-known meat spoiler *Brochothrix thermosphacta* in broiler meat [8, 33]. Our study, however, found *B. thermosphacta* to make up only a minute portion of the active spoilage microbiome. This is likely to result from the anaerobic packaging conditions that gave LAB an advantage to outcompete *B. thermosphacta* almost completely, based on the microbiome analysis results, fungi could not compete with bacteria in the spoilage microbiome of anoxic MA packaged broiler meat. They were detected in the beginning but the reads aligning to them quickly started falling when the broiler meat aged (Figure 2). Fungi have been known to take part in beef spoilage [11], but to keep its red colour, beef is packaged under high oxygen MAP, improving the conditions for the growth and survival of fungi.

## 5. Conclusions

Meat spoilage is based on complex interspecies interactions in the course of time, and our findings fill one of the many gaps in knowledge of the spoilage process. This lack of knowledge related to meat microbiomes has also been recently pointed out by Wang *et al.* [26]. A persistent difficulty in food spoilage research is that the concentration of bacteria in MA packaged meat does not correlate linearly with sensory spoilage changes. Our research demonstrates the potential of RNA-seq coupled with metabolomics in better understanding the underlying processes of meat spoilage and studying active microbiomes in general. We were able to detect *V. proximus* within the more abundant species in the active microbiome and the change it had in the pathway activity distribution. Future studies should be targeted to increase understanding of the role *Vagococcus* as a food spoiler.

Traditional culturing or 16S rRNA amplicon sequencing overlook the phenomena that lay under the main frame of microbiomes but may provide interesting insights in understanding meat spoilage better.

Altogether, our results illustrate how the microbial and functional changes underpin the chemical and sensory spoilage of broiler meat and demonstrate the value of refrigeration temperatures below 6°C. This study also demonstrates the benefit of studying the transcriptional activity of the whole microbiome to uncover the underlying dynamics instead of only looking at the most abundant and culturable species.

## Declarations

### Ethics approval and consent to participate

The broiler chicken used in this study was produced by a large-scale commercial meat producer and intended for human consumption. No ethical approval was required for this study.

### Consent for publication

Not applicable.

### Availability of data and materials

The raw RNA-seq data for both temperatures is available in the SRA archive of NCBI under accessions SRR38506858-SRR38506902 of BioProject PRJNA1464459.

All scripts and code used in this article can be found at GitHub

### Authors’ contributions

JM processed the raw RNA-seq data, analysed and interpreted the active microbiome composition and was a major contributor in writing the manuscript. EN analysed the microbiome pathway activity over time. EJ performed and interpreted the results of the non-targeted metabolomics. PJ supervised and assisted with the RNA-seq data analysis. JB obtained the funding for this research, supervised and assisted with the interpretation of the results.

## Acknowledgements

We would like to thank Henna Niinivirta for her skilful work in the laboratory, without which this research would not have been possible.

