## Supplementary_figures for "Longitudinal RNA Seq analyses reveal the prominent role of *Vagococcus* in broiler meat spoilage microbiome"

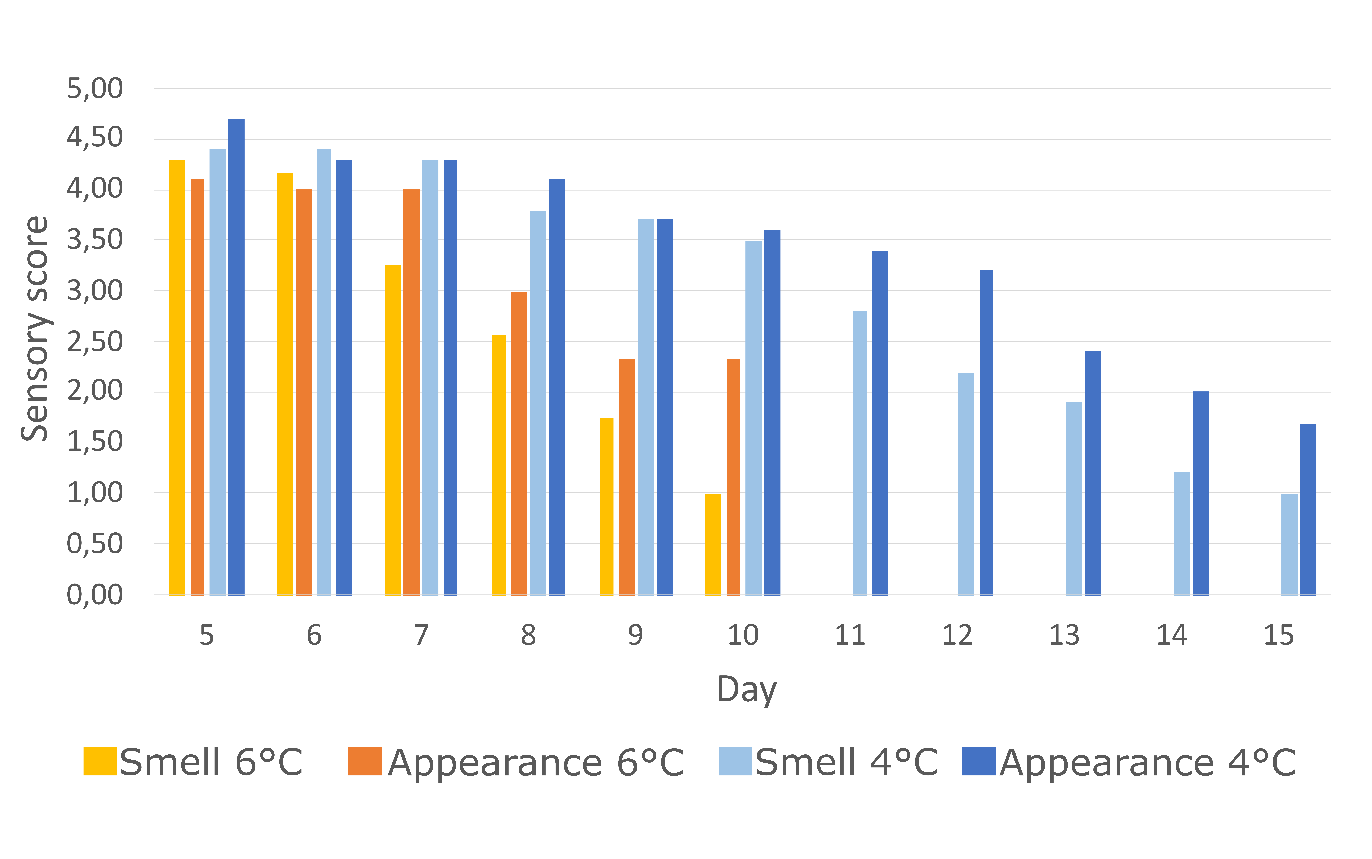

Supplementary Figure 1. Sensory scores of broiler meat at 4°C and 6°C.

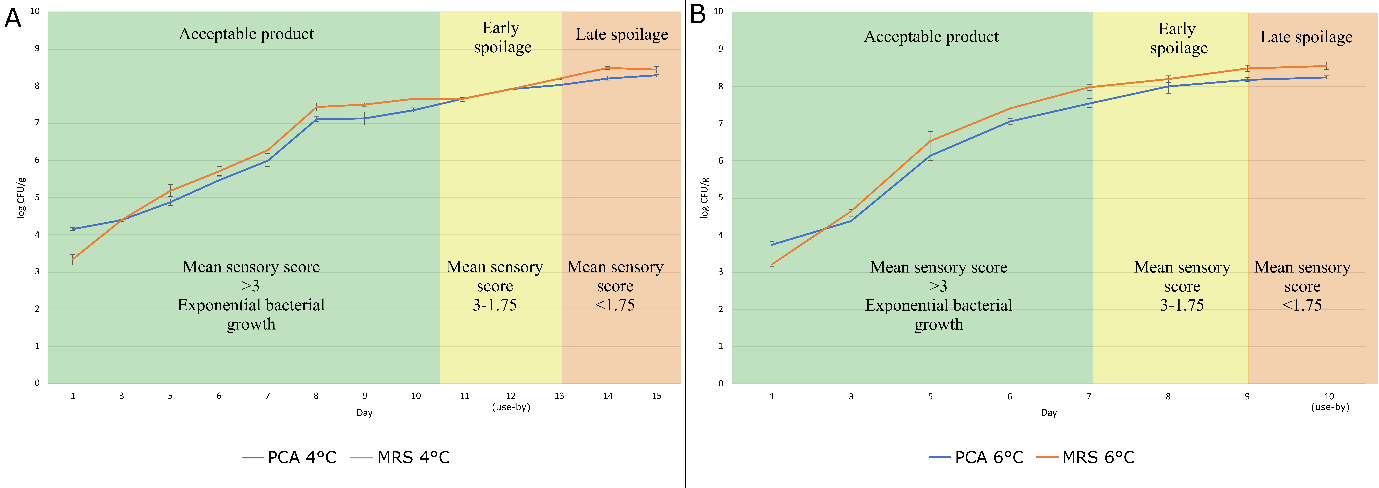

Supplementary Figure 2. Bacterial growth of the broiler samples on MRS and PCA combined with the phase of spoilage according to the sensory scores.

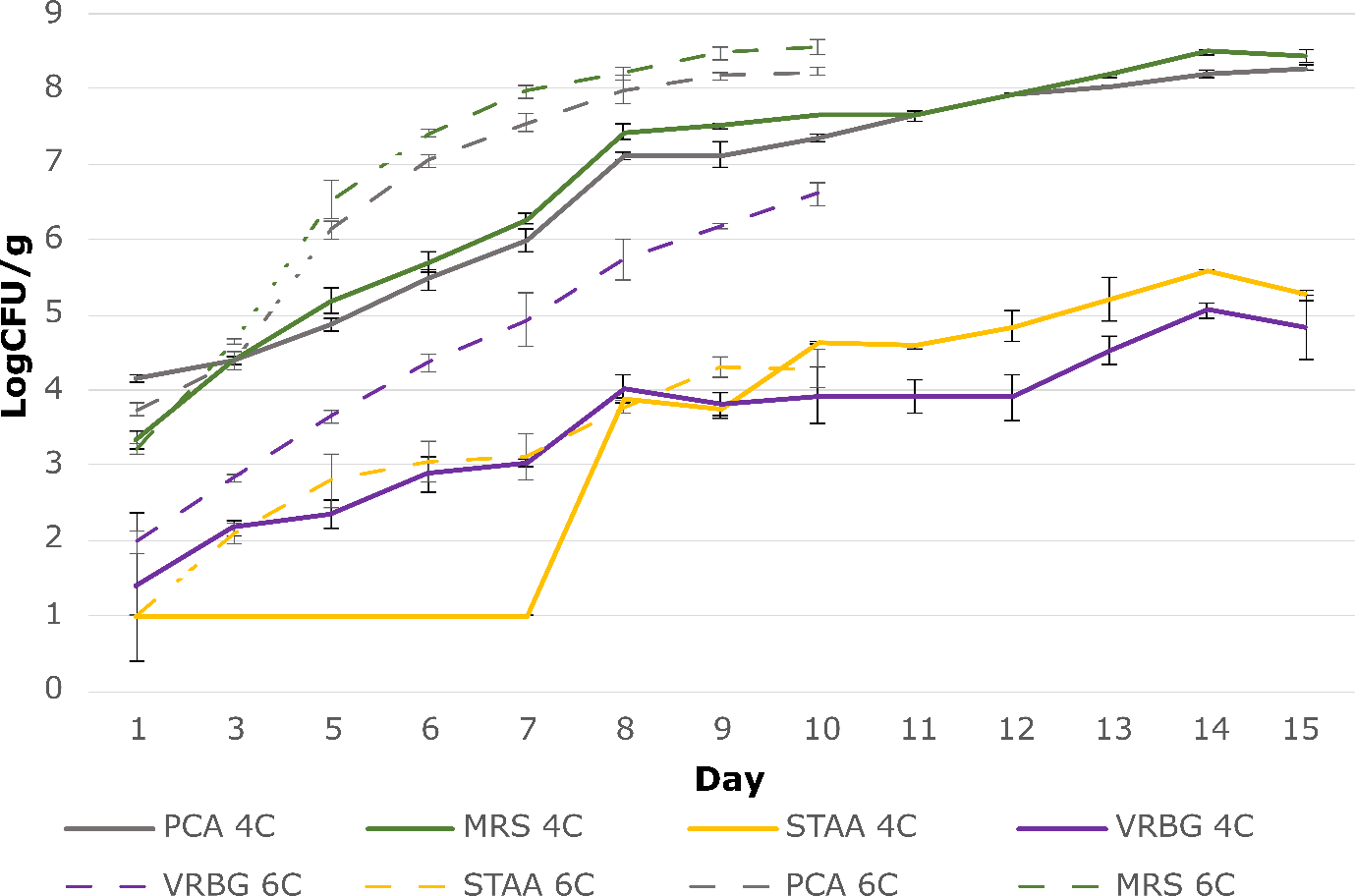

Supplementary Figure 3. Bacterial growth of the broiler samples on four different media at both 4°C and 6°C.

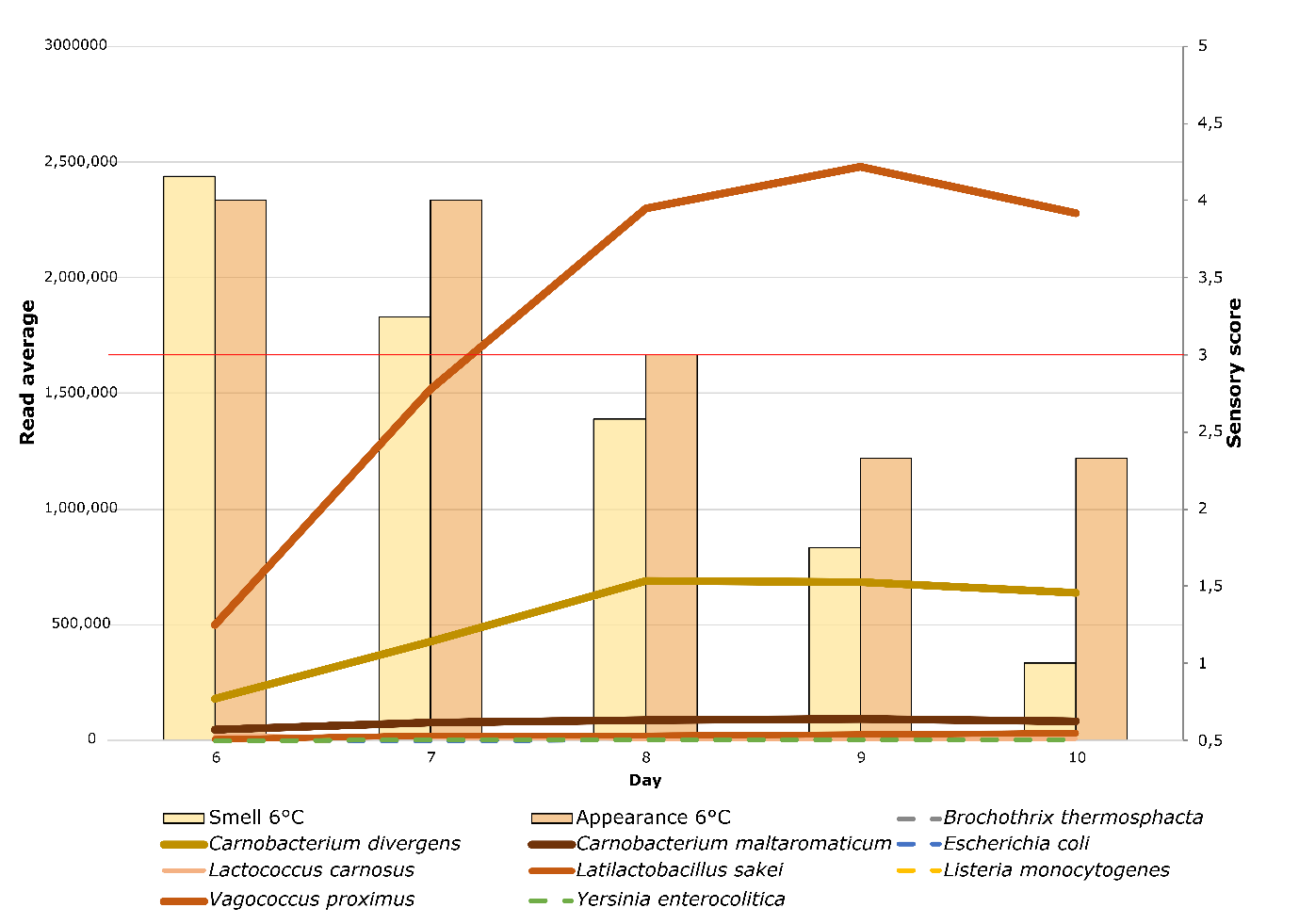

Supplementary Figure 4. Average read counts of the most prominent bacteria in the active spoilage microbiome at 6°C.

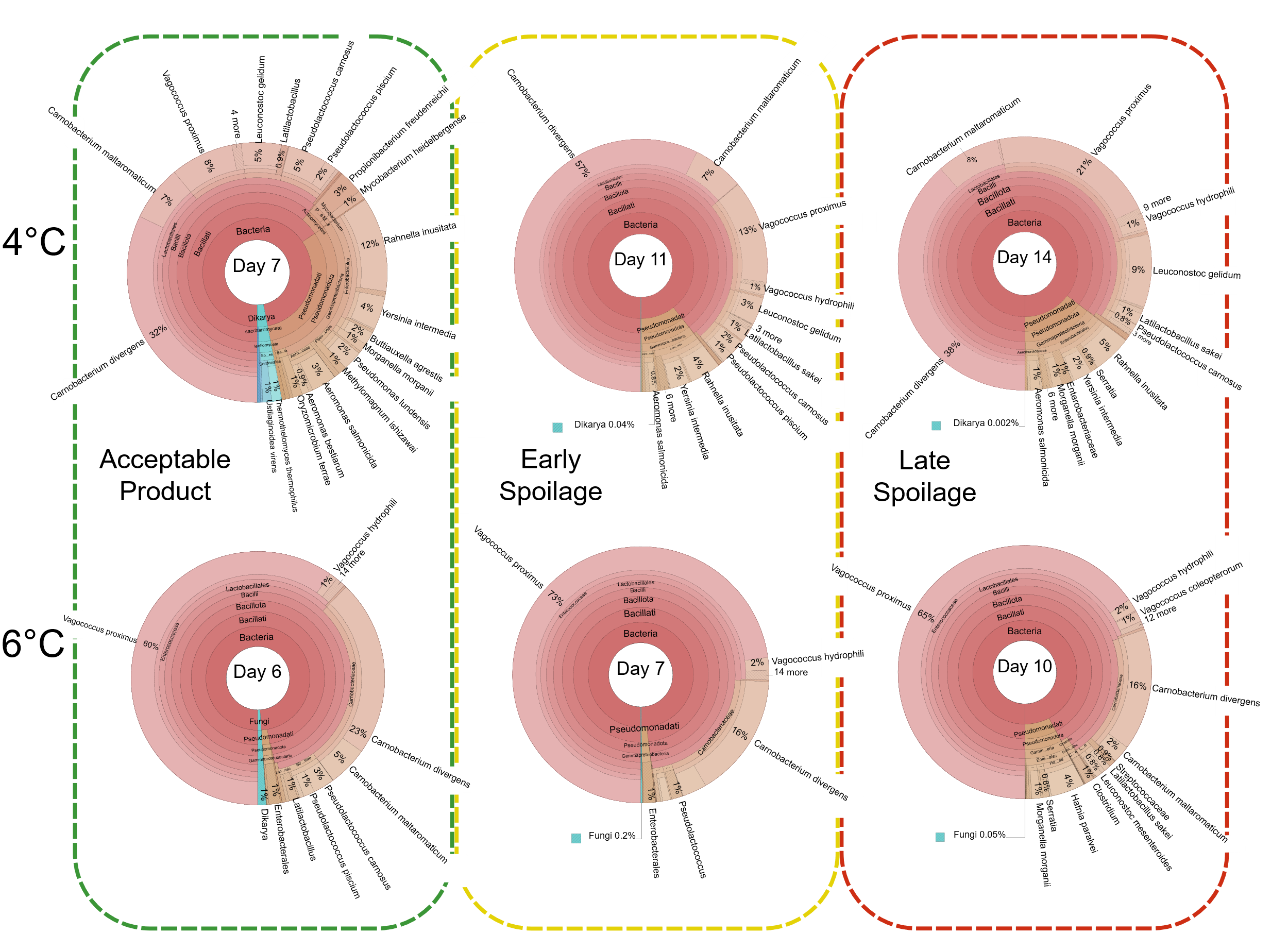

Supplementary Figure 5. Comparison of the microbial communities during different phases of spoilage between the two temperatures.
